# Genomics of viruses infecting green and purple sulfur bacteria in two euxinic lakes

**DOI:** 10.1101/2022.07.26.501573

**Authors:** P. J. Hesketh-Best, A. Bosco-Santos, S. L. Garcia, J. P. Werne, W. P. Gilhooly, C. B. Silveira

## Abstract

Viral infections of marine bacteria modulate the rates of primary production and the cycling of organic and inorganic matter in the world’s oceans. Here, we investigated the hypothesis that viral infections influence the ecology of purple and green sulfur bacteria (PSB and GSB) in anoxic and sulfidic (euxinic) lakes, modern analogs of early Earth oceans. Over 200 high and medium quality viral contigs were identified in long-read metagenomes from the sediments and water column of Lime Blue and Poison Lake, respectively. We compared these sequences with 94 predicted prophages identified in the complete genomes of PSB (n = 213) and GSB (n = 33). Viral genomes carrying *psbA*, encoding the small subunit of photosystem II protein, were present in all three datasets (sediment, water column, and complete genomes). The ubiquity of these genes suggests that PSB and GSB viruses interfere with the light reactions of sulfur-oxidizing autotrophs in a process similar to viral modulation of photosynthesis in Cyanobacteria. Viruses predicted to infect PSB and GSB also encoded auxiliary metabolic genes involved in reductive sulfur assimilation as cysteine, a pathway not yet described in these sulfur bacteria, as well as genes involved in pigment production (*crtF*) and carbon fixation (CP12, *zwf*, PGD). These observations highlight the potential for viral modulation of metabolic markers used as proxies to interpret biogeochemical processes in early Earth oceans.

## Introduction

Before the Great Oxygenation Event, culminating 2.33 billion years ago, anoxygenic prototrophic bacteria catalyzed most marine primary production, influencing ocean stratification and the planet’s oxidant balance (Kappler and Straub, 2005; Johnston *et al.*, 2009; Farquhar *et al.*, 2011). Our view of the biological processes mediated by microorganisms in ancient Earth has largely been informed by the assumption that there is a direct feedback between environmental gradients (such as pH, salinity, temperature, nutrients, oxygen/sulfide, carbon source, sulfate availability) and microbial community composition and distribution (Borda et al., 2001; Brocks et al., 2005; Kappler and Straub, 2005; Brocks and Schaeffer, 2008; Johnston et al., 2009; Farquhar et al., 2011; Fakhraee et al., 2019; Ozaki et al., 2019; Bosco-Santos et al., 2020). While these assumptions may be broadly correct, they produce an aggregate view of the complexity of microbial interactions that overlooks a critical regulator of pre-GOE biogeochemical cycles. Here we propose that a largely unexplored biotic factor controls the distribution and activity of anoxygenic sulfide oxidizing phototrophs: viral infection.

Anoxygenic sulfide oxidizing phototrophs from green (GSB, family *Chlorobiaceae*; Garrity et al., 2001) and purple (PSB, families *Chromatiaceae* and *Ectothiorhodospiraceae*; Imhoff, 2003) sulfur bacteria inhabit the euxinic photic zone, where sulfide intercepts the sunlit portions of stratified marine and lacustrine anoxic water columns. These primary producer groups have narrow environmental optimal requirements as micro-oxic to anoxic conditions, free sulfide, and sunlight, with GSB more adapted to lower light levels and PSB more tolerant to dissolved oxygen (Hamilton *et al.*, 2014). Consequently, GSB and PSB light-harvesting pigments and biomarkers are potential proxies for diagnosing basin depth and redox state in the geologic record, providing clues about past biological processes and environmental conditions (Koopmans *et al.*, 1996; Brocks and Schaeffer, 2008; Brocks and Banfield, 2009). In other words, the presence or preservation (as diagenetic products) of GSB (chlorobactene and isorenieratene) and PSB (okenone) biomarkers are interpreted as deep or shallow redoxcline, respectively (Brocks *et al.*, 2005; Brocks and Schaeffer, 2008). Yet, a growing body of evidence shows that the distribution of GSB and PSB in modern water columns is not as tightly correlated to physical and chemical conditions (e.g., sulfide and light) as previously thought, suggesting that biological interactions play a significant role in defining the distribution of these phototrophs (Massé *et al.*, 2002; Hamilton *et al.*, 2014; Llorens-Marès *et al.*, 2017).

Bacteriophages, also known as phages, are viruses that infect bacteria and can laterally transfer genes, modulate gene expression, and control host population dynamics (Breitbart, 2012). In the modern surface ocean, viral predation is responsible for the daily turnover of about 25% of the bacterioplankton (Breitbart *et al.*, 2018). Viruses of modern Cyanobacteria encode genes for enzymes in the Calvin cycle, blocking carbon fixation during infection while increasing nucleotide production through the Pentose Phosphate pathway (Thompson *et al.*, 2011). Most of these carbon metabolism pathways are shared between Cyanobacteria and sulfide oxidizing phototrophs, and viral interference with carbon fixation in GSB and PSB is possible. A recent study showed that lake GSB populations were concurrently infected with 2-8 viruses per cell (Berg *et al.*, 2021). One GSB host was consistently associated with two prophages with a nearly 100% infection rate for over 10 years (Berg *et al.*, 2021). GSB genomes have high signatures of horizontal gene transfer, reaching 24% of all genes in *Chlorobaculum tepidum* (Nakamura *et al.*, 2004). Likewise, phages infecting oxygenic phototrophs encode many genes involved in the synthesis of light harvesting pigments (*hol, pebS, cpeT, pcyA*) (Breitbart *et al.*, 2018). Therefore, probing phage regulation of pigment synthesis in anoxygenic phototrophs is a necessary step toward understanding the ecology of biomarker production in euxinic systems, with implications for interpreting the deep time record based on diagenetic products of pigment biomarkers.

Here, we identify through long-read metagenomic sequencing the genomes of viruses putatively infecting GSB and PSB inhabiting euxinic lakes. These viruses encode several genes involved in carbon fixation, sulfur metabolism, and pigment production. Based on these observations, we propose that GSB and PSB viruses manipulate host metabolism, potentially influencing these autotrophs’ biogeochemical signatures in the geologic record.

## Methods

### Sampling

Poison Lake water column (2L) was collected from a boat using a peristaltic pump to obtain a sample from the sulfidic zone. Subsamples (50ml) were immediately frozen until further laboratory processing. In the laboratory, samples were defrosted and incubated overnight at 4 °C with Polyethylene Glycol 8000 10 %. Samples were centrifuged at 5000 g for 2 hours at 4 °C and the pellet was collected for DNA extraction with a DNeasy PowerSoil kit (Qiagen, Germany). The sediment from Lime Blue was collected with a freeze core (modified from Stocker and Williams, 1972). The sediments were sectioned within a sterile flow bench to prevent organic contamination. An archive section (~1/3 of the core’s width) was preserved and stored at −80°C for future use. Sediment was collected every 2cm, for a total of 25 samples dating back to the deposition year of 1424. Sediment subsamples (1g) were extracted using the DNeasy PowerSoil kit (Qiagen, Germany), following manufacturer’s instructions. Preliminary 16S sequencing of these samples revealed an abundance of anoxygenic photosynthetic bacteria in the sedimentary record of Lime Blue in the last ~580 years where it is possible to observe that above 15 cm deep the relative abundance of each family increased more than 100%. The top 2cm sample was sequenced here.

### Long-read sequencing

Metagenomic libraries were prepared using the ONT Ligation Sequencing Kit (SKQ- LSK110, Oxford Nanopore Technologies) following the manufacturer’s instructions. In short, 1mg of dsDNA was End-prepped and repaired to ligate a poly-A tail using the NEBNext Companion Module for Oxford Nanopore Technologies Ligation Sequencing (cat # E7180S), before sequencing adaptors were ligated onto the ends. Between each step, DNA was cleaned using 1.8X Agencourt AMPure XP beads (Beckman, USA), washing the beads with 70% molecular grade Ethyl alcohol (Sigma-Aldrich, USA) before suspending in Nuclease-free water (Fisher, USA). Sequencing libraries were loaded onto and sequenced using a FLO-MINSP6 flow cell, and sequencing protocol was run for 48 hrs.

### Identification of viruses in metagenomes and publicly available PSB and GSB genomes

ONT sequencing adaptors were trimmed using Porechop v0.2.4 (https://github.com/rrwick/Porechop), and trimmed reads were assembled with Flye v2.9 (Lin *et al.*, 2016; Kolmogorov *et al.*, 2020) using the *--meta* parameter. In parallel, low quality and short reads were removed by NanoFilt v2.6.0 (De Coster *et al.*, 2018) to a minimum Q-value of 9 and length of 1 kb. Both the metaFlye contigs and quality filtered ONT reads were utilized for the detection of phages by VIBRANT v1.2.1, a bioinformatics pipeline using Hidden Markov Model (HMM) searches to identify clusters of viral genes in unknown sequences, allowing the sorting of high-confidence viral genomes and genome fragments within complex samples (Kieft *et al.*, 2020).

Publicly available bacterial genomes with a completion level of ‘complete genome’, ‘scaffold’ and ‘contig’ belonging to the two PSB families *Chromatiaceae* (98 genomes) and *Ectothiorhodospiraceae* (115 genomes), and the GSB phyla Chlorobiota (33 genomes) were retrieved from NCBI in 2022 (Supplementary Table 1). Putative prophages were identified in these genomes using VIBRANT v1.2.1.

The viral genomes and genome fragments were screened for the presence of carbon, sulfur, and pigment-related auxiliary metabolic genes (AMGs) and their potential for lysogeny (presence of transposases and integrases) through HMM comparisons with three databases: Pfam, VOGs, and SEED. Viral genomes containing AMGs of interest were visualized using the R package genoPlotR v0.8.11 (Guy *et al.*, 2010). For a small selection of phages containing AMGs of interest, the Max Planck Institute (MPI) HHpred server (Zimmermann *et al.*, 2018) was utilized to improve genome annotations (E-value <0.01 and Probability > 80%), in addition to the Phage Artificial Neural Networks (PhANNs, Cantu et al., 2020) to confirm phage structural proteins (Confidence > 80%). To analyze the abundance and coverage of these putative viral genomes in the environment, trimmed reads were mapped to the viral contig database at high stringency (>95% identity). An outlined summary of the entire workflow can be seen in Supplementary Figure 1.

### Generation and quality control of MAGs

Metagenome-assembled genomes (MAGs) of Bacteria were generated by mapping raw ONT reads to metaFlye contigs with Minimap2 v2.24 (Li, 2018). Subsequence .SAM files were compressed, sorted, and indexed with samtools v1.9 (Danecek *et al.*, 2021). Metagenomic bins were generated using three binning programs: CONCOCT v1.0 (Alneberg *et al.*, 2014), MetaBAT2 v2.12.1 (Kang *et al.*, 2019), and MaxBin2 v2.2.6 (Wu *et al.*, 2016). Resulting bins were refined using MetaWRAP v1.3 *bin_refinement* module (Uritskiy *et al.*, 2018), and refined bins were assessed for contamination and completion with CheckM v1.2.0 (Parks *et al.*, 2015). Bins with ≥ 50% completion and ≤ 10% contamination were kept for further analyses. MAG depth of coverage (mean) was quantified by mapping clean reads to the metagenomic bins and taking the mean percentage of reads mapped. ONT reads and contigs were taxonomically classified by Kraken v2.0 (Wood and Salzberg, 2014; Wood *et al.*, 2019) and abundance estimated by Bracken (Bayesian Re-estimation of Abundance after Classification with KrakEN) v2.7 (Lu *et al.*, 2017).

### Phylogeny and taxonomic classification of MAGs

The 16S rRNA gene, if present, was extracted from MAGs and inspected for contamination directly from the genomes and metagenomic bins using ContEst16S. Neighbours for MAG 16Sr rRNA gene were determined using BLASTn against the NCBI database, and when an uncultured clone was the only match, additional BLASTn of the 16S rRNA was run against the SILVA database. Sequences were aligned using MAFFT v7.5 (Katoh *et al.*, 2002; Katoh and Standley, 2013), and a maximum-likelihood tree was generated using RAxML-NG v0.9 at 200 bootstraps (Kozlov *et al.*, 2019). The tree was visualized and edited for readability using the interactive tree of life (iTOL) v6 (Letunic and Bork, 2007, 2021).

### Phage host prediction

Viral genomes observed within bacterial genome fragments were identified as lysogenic. Viral hosts were identified using a combination of gene homologies, the presence of tRNAs, and CRISPR (clustered regularly interspaced short palindromic repeats) spacers (Coutinho *et al.*, 2017; Borges *et al.*, 2022). (I) Sequence homology matches were made from the phages identified from Lime Blue and Poison Lake to databases generated from PSB/GSB genomes retrieved from NCBI, and MAGs generated in this study using BLASTn (Camacho *et al.*, 2009). Only hits >80% sequence identity across a minimum alignment of 1,000 nucleotides were considered as possible hosts for NCBI and RefSeq genomes, and 95% sequence identity against MAGs. (II) A database was created with the CRISPR spacers from PSB, GSB genomes and MAG using minCED v0.4.3 (Mining CRISPRs in Environmental Datasets; https://github.com/ctSkennerton/minced), which uses CRISPR Recognition Tools (CRT) v1.2 (Bland *et al.*, 2007), and sequence homology matches were made against the phages using BLASTn with the parameter *-task “blastn-short”*, hits were only considered with a maximum of 2 mismatches/gaps, 100% sequenced identity, and minimum length of 20 nucleotides. (III) Phage tRNAs were detected using tRNAScan-SE v2.0 (Lowe and Chan, 2016), and matched against PSB/GSB/MAG genomes database for hits using BLASTn, a confident hit was considered at ≥ 90% sequence identity and ≥ 90% coverage. (IV) Phylogenomic analysis was performed against the GL-UVAB (Gene Lineage of Uncultured Viruses of Archaea and Bacteria) reference database, as previously described (Coutinho *et al.*, 2019).

## Results

### Bacterial community composition

Nanopore sequencing generated 3.9 x10^6^ reads from Lime Blue (LB) sediment and 19.2 x10^6^ reads from Poison Lake (PL) water. Trimming, quality filtering short (reads ≤ 1000 bp) and low-quality reads (*Q-value* < 9) removed 96 and 93% of reads from LB and PL, respectively. Assemblies generated 40,807 LB contigs and 4,310 PL contigs. A broader GSB and PSB bacterial diversity was identified by taxonomic classification of reads than by metagenomic binning (Supplementary Figure 2-3). According to Kraken2 classification, clean reads from LB and PL were dominated by members of the phylum Proteobacteria (reads: 48.77% LB and 70.51% PL; contigs: 45.11% LB and 59.31% PL) (Figure 1A), of which Gammaproteobacteria was the most abundant class for both. For LB sediment, at read level, LB was dominated by the anaerobic specialist order Enterobacterales (22.50%), but this was not reflected post assembly, with only 5.89% of contigs classified as Enterobacterales. For PL, the order of phototrophic sulfur bacteria Chromatiales was the most abundant Gammaproteobacteria (reads: 11.38%; contigs: 8.60%).

**Figure 1.**
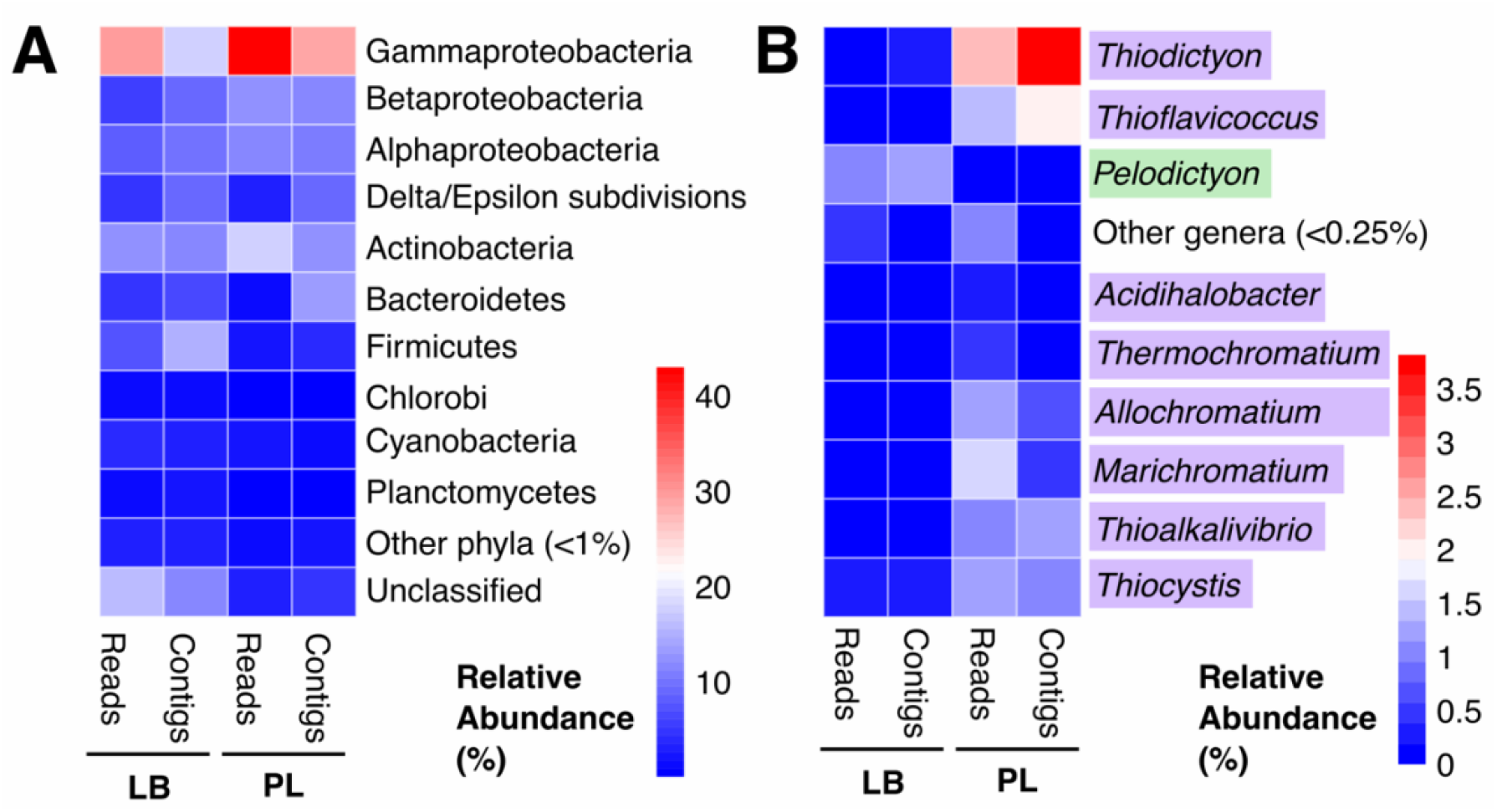
Phylum and genus level taxonomic classification of reads and contigs from Lime Blue (LB) and Poison Lake (PL). (A) Phylum level read and contig diversity with members of the phylum Proteobacteria split into class level, and (B) genus level diversity of Chlorobi, *Chromatiaceae* and *Ectothiorhodospiraceae*, with PSB highlighted in purple and GSB highlighted in green.

Within the order Chromatiales, PL water samples show higher relative abundances of families *Chromatiaceae* (reads: 8.45%; contigs: 8.38%) and *Ectothiorhodospiraceae* (reads: 1.98%; contigs: 1.25%) and contained a variety of PSB genera in abundances ranging from <0.25% to 3.82%, with *Thiodictyon* spp. (reads: 2.34%; contigs: 3.82%) being the most abundant. In contrast, phototrophic sulfur bacteria represented a smaller fraction of the metagenomic dataset in LB sediment, with a greater abundance of GSB from phylum Chlorobi (reads: 1.26%; contigs: 1.24%) than PSB, order Chromatiales (reads: 0.83%; contigs: 1.05%). The genera *Pelodictyion* spp. was the most abundant GSB (reads: 0.10%; contigs: 1.15%), and *Thiocystis* spp. (reads: 0.25%; contigs: 0.36%) was the most abundant PSB.

### Bacterial MAG recovery and phylogeny

Bacterial genomes were binned from LB sediment and PL water assemblies, resulting in 17 MAGs with a minimum completion of 50% and maximum contamination of 10% (Table 1). Of these MAGs, seven were classified as high quality (≥70% completion, ≤ 10% contamination). The most abundant MAG, as measured by mean coverage of Nanopore reads mapped to the bins, was the PL_bin01 (mean = 2.74%), classified as a potential PSB belonging to the genera *Thiohalocapsa*, and LB_bin03 (mean = 0.11%), classified as Candidatus Komeilibacteria (Supp. Figure 2).

**Table 1.** Taxonomy and quality of MAGs. Quality was estimated by CheckM and taxonomy by 16S rRNA gene using the SILVA ACT service. Values above the threshold for a high-quality MAG are denoted in bold (≥ 70% completion, and ≤ 10% contamination). (*) denotes potential PSB MAG, and (†) denotes a MAG identified as potential host to a phage by CRISPR spacers. (Comp., bin completeness; Contam., bin contamination; Strain het., proportion of the contamination that originates from the same or similar strains)

| Bin | CheckM Marker lineage | 16S rRNA placement base on the SILVA-ACT |  | Comp. | Contam. | Strain het. | Mean cov. (%) |
| --- | --- | --- | --- | --- | --- | --- | --- |
|  |  | Phylum | Lowest taxonomic classification |  |  |  |  |
| LB_bin01 | k_Bacteria (UID2569) | Spirochaetota | g_Treponema | 60.12 | 1.10 | 0.00 | 0.009 |
| LB_bin02 | k_Bacteria (UID2569) | Latescibacterota | p_Latescibacterota | 58.63 | 1.10 | 0.00 | 0.016 |
| LB_bin03 | k_Bacteria (UID2569) | Patescibacteria | o_Candidatus Komeilibacteria | 62.23 | 0.00 | 0.00 | 0.106 |
| LB_bin04 | k_Bacteria (UID2569) | Latescibacterota | o_WCHB1-41 | 55.01 | 1.10 | 0.00 | 0.011 |
| LB_bin05 | k_Bacteria (UID2569) | No 16S rRNA gene detected |  | 70.8 | 4.65 | 33.33 | 0.015 |
| LB_bin06 | k_Bacteria (UID2569) | Verrucomicrobiota | p_Verrucomicrobiota | 36.55 | 3.49 | 0.00 | 0.008 |
| LB_bin07 | k_Bacteria (UID2569) | Chloroflexi | o_MSBL5 (contaminated) | 74.72 | 7.59 | 20.0 | 0.005 |
| LB_bin08 | k_Bacteria (UID2569) | No 16S rRNA gene detected |  | 61.32 | 2.63 | 16.67 | 0.018 |
| LB_bin09 | k_Bacteria (UID2569) | k_Archaea | f_Candidatus Iainarchaeum | 56.63 | 0.00 | 0.00 | 0.005 |
| LB_bin10 | k_Bacteria (UID2569) | Bacteroidota | g_Ignavibacterium | 58.8 | 3.03 | 28.57 | 0.012 |
| LB_bin11 | p_Bacteroidetes (UID2605) | No 16S rRNA gene detected |  | 59.05 | 9.14 | 0.00 | 0.005 |
| PL_bin01*† | c_Gammaproteobacteria (UID4274) | Proteobacteria | g_Thiohalocapsa (Chromatiales) | 85.17 | 1.17 | 40.0 | 2.724 |
| PL_bin02 | k_Bacteria (UID2569) | Latescibacterota | o_WCHB1-41 | 76.74 | 3.05 | 9.09 | 0.009 |
| PL_bin03 | k_Bacteria (UID2569) | Deinococcota | g_Truepera | 76.11 | 2.97 | 71.43 | 0.010 |
| PL_bin04 | c_Deltaproteobacteria (UID3217) | Desulfobacterota | g_Desuloanatronum | 89.53 | 0.60 | 100 | 0.016 |
| PL_bin05 | k_Bacteria (UID2569) | Planctomycetota | f_KCLunmb-38-53 | 82.29 | 3.42 | 11.11 | 0.027 |
| PL_bin06 | k_Bacteria (UID2569) | No 16S rRNA gene detected |  | 68.8 | 0.00 | 0.00 | 0.01 |

### Diversity of PSB-infecting phages

From publicly available PSB and GSB genomes, VIBRANT identified a total of 32 phages of high quality (HQ), 36 of medium quality (MQ) and 183 of low quality (LQ). Of the HQ and MQ phages, 64 were from Chromatiales genomes (33 *Chromatidales* phages, and 31 *Ectothiorhodospiraceae*). The majority (63) of HQ and MQ phages were classified as lysogenic, and of the eight phages classified as lytic, three were complete/circular. No Chlorobi phages were identified as lysogenic, indicating the absence of known integration enzymes in these integrated prophages identified within their hosts’ genomes. In total, four complete genomes were predicted, one from the GSB *Chlorobium limicola* strain Frasassi, one from *Thiocystis violacea* strain DSM 207, and two from *Thiohalocapsa* sp. ML1 and *Halochromatium roseum* DSM 18859.

From the metagenomic sequences, VBRANT identified 2,742 phages from LB contigs (100 MQ phages and 24 HQ phages). From PL metagenomic reads, 5,806 phages were identified, all of which were LQ phages. Two phages were classified as complete and circular from LB and none from PL. Contigs did not improve the quality of predicted phages in PL, and filtered PL reads were utilized for further viral analyses.

Bacterial MAGs and publicly available PSB and GSB genomes were utilized to predict hosts of PSB- and GSB-infecting phages using a combination of BLASTn against PSB and GSB genomes, CRISPR-spacers of the putative host and, if available, tRNA sequences (Supp. Table 2). Homology matches against a database of PSB/GSB genomes predicted hosts for 5,451 phages (12 LB phages, and 5,439 PL), with the most common host for phages from both samples being *Chromatium weissei* DSM 5161. Homology matches against MAGs resulted in 547 high confidence predictions, with the PSB PL_bin01 (*Thiohalocapsa* sp.) and the PL_bin04 (*Desuloanatronum* sp.) as the most common predicted phage hosts. High confidence phage-host linkages based on CRISPR-spacer homology matches with 100% identity, and > 20 nucleotide coverage predicted hosts for 54 phages (44 LB phages, and 10 PL phages). The most common host for LB phages was *Ectothiorhodospira* spp., while for PL phages predicted hosts included *Allochromatium* spp*., Chlorobium* spp. and *Thiohalocapsa* spp. Homology matches to a database of tRNA sequences only yielded four predictions, with *Thiohalocapsa* sp. ML1 being the only predicted hosts for three PL phages, and *Thiorhodovibrio winogradskyi* strain 6511 for one LB phage.

The LB and PL phages were compared to reference viral genomes using clustering based on gene sharing distance (Coutinho *et al.*, 2019) (Figure 2). Most LB phages and PSB and GSB phage clusters had long branch lengths, evidence of low similarity between phages predicted by VIBRANT in this study and the reference viral genomes (Supplementary Figure 4). Several clusters were formed exclusively of LB phages. Only one cluster of LB phages closely relates to a predicted phage from PSB genomes, despite clustering with reference GL-UVAB phages with PSB hosts. This may indicate that many of the phages detected in this study infect uncharacterized bacterial hosts. The GL-UVAB viruses related to the viruses identified here infected *Chromatidales* and *Ectothiorhodospiraceae*, with the taxonomy of most hosts unresolved beyond family level, including viruses of *Thioalkalivibrio*, *Thiorhodoccocus*, and *Thiorhodovibrio*.

**Figure 2.**
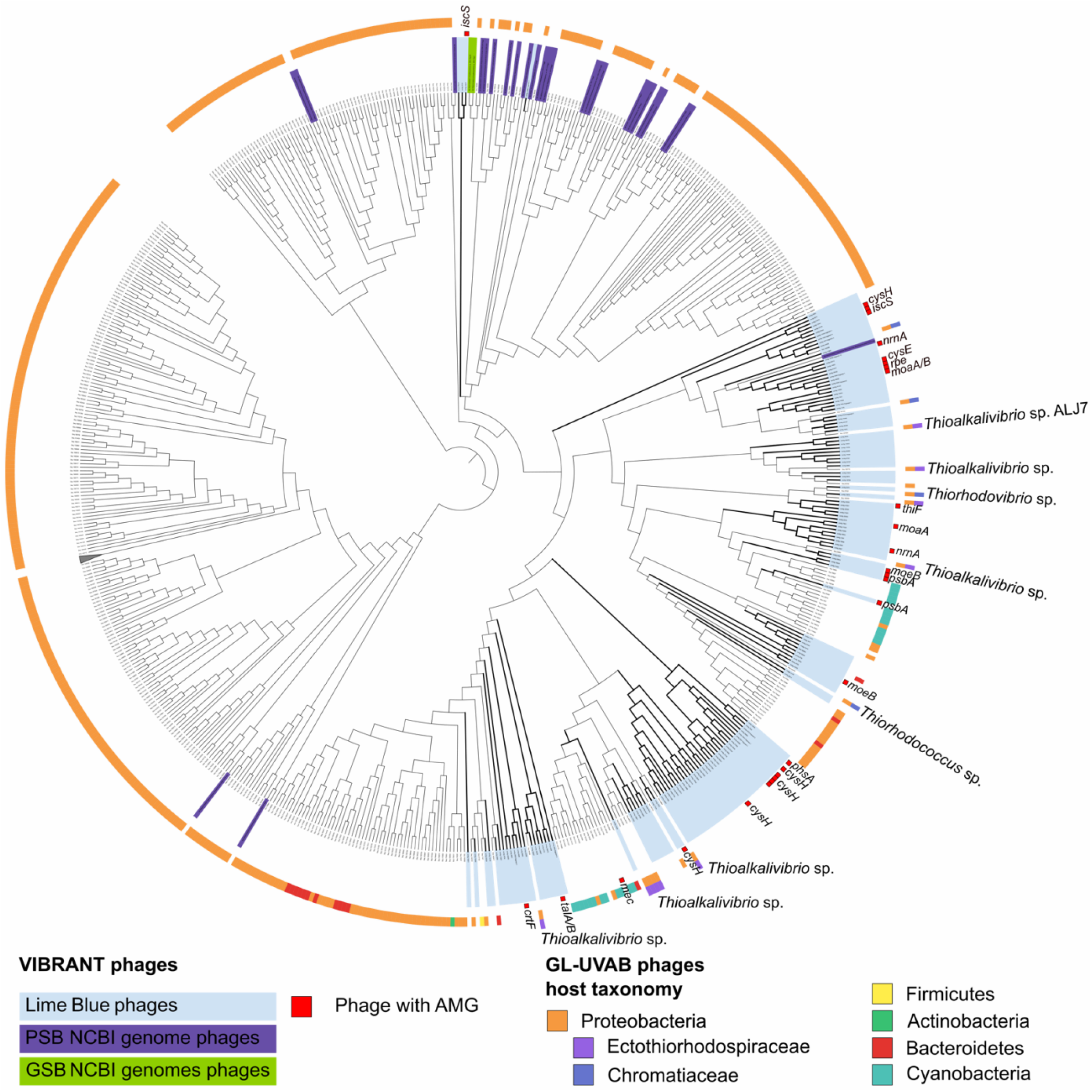
Clustering of VIBRANT phages from LB metagenome (blue) and PSB genomes (purple) and GSB genomes (green) and the reference phage genomes based on their gene-sharing Dice distances. The VIBRANT phages from Lime Blue, many of which contained AMGs of interest (inner ring), form novel branches with low similarity to reference phage genomes. Where known, the host genera of the reference PSB infecting phage were listed. The branch lengths are ignored to better display clustering topology, for a version displaying branch length, see Supplementary Figure 4.

### Phage AMGs influencing diverse metabolic pathways

A total of 52 and 96 AMGs were detected from PL and LB phages respectively, with the AMGs representing 153 distinct KEGG pathways, including photosynthesis, sulfur metabolism and relay, pigment synthesis, Calvin Cycle, and Pentose Phosphate Pathway (PPP) (Figure 3A). Five phages from the *Chromatidales* genomes contained AMGs involved in sulfur metabolism and relay (*cysH, moeB*, and *mec*). The bacterial hosts of these phages included *C. weisse* DSM 5161 (*cysH* and *mec*), *T. violacea* DSM 207 (*cysH), Thiospirillum jenense* DSM 216 (*moeB*), and *Allochromatium humboldtianum* DSM 21881 (*mec*). The phages predicted from *T. jenense* and *A. humboldtianum* encoding AMGs were classified as lysogenic. A single lysogenic phage from an *Ectothiorhodospirales* genome contained a *cysH* that was detected from the plasmid pTK9001 of *Thioalkalivibrio* sp. K90mix. A single lytic phage from the GSB *Chlorobium limicola* strain Frasassi contained the CP12 gene involved in blocking carbon fixation through the Calvin Cycle in Cyanobacteria.

**Figure 3.**
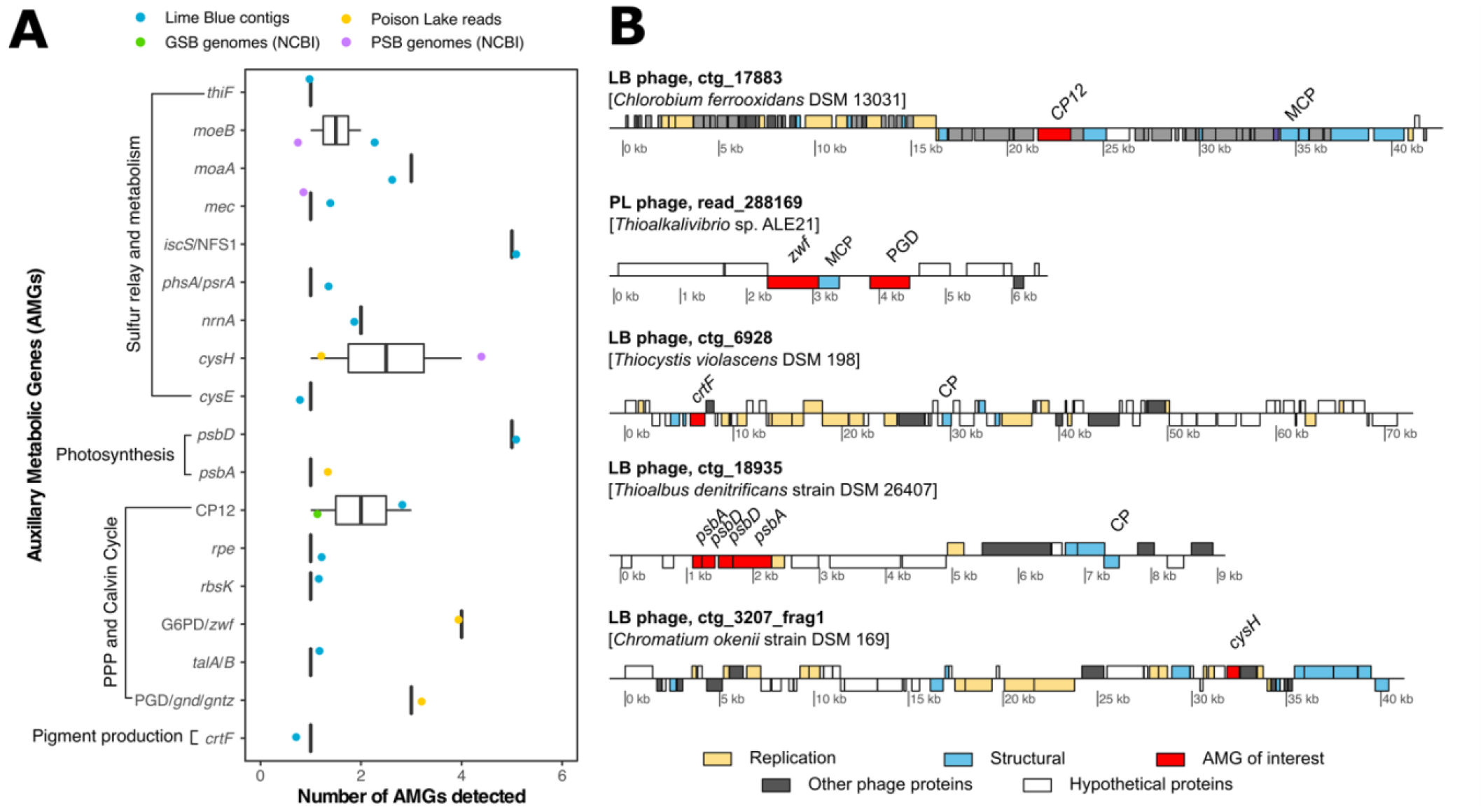
(A) AMG abundances and (B) genome maps of putative phages containing AMGs of interest. AMGs were identified by VIBRANT except for CP12, which was identified by BLASTp of phage ORFs to a database of available CP12 proteins from UniProt. Putative hosts identified based on CRISPR spacers are indicated for each phage (LB, Lime Blue; PL, Poison Lake; MCP; major capsid protein; CP; putative capsid protein)

LB phages encoded several AMGs involved in sulfur metabolism (*cysE*, *nrnA* and *pshA*) and sulfur relay (*moeB*, *thiF* and *iscS*). While most of the AMGs were detected in phages predicted to be lytic, four LB lysogenic phages contained a copy of *cysH, moeA*, and *nrnA.* No PL phages were observed to contain sulfur metabolism or relay AMGs. AMGs related to photosynthesis (*psbA* and *psbD*) were present in both PL and LB lytic phages, with four phages containing multiple copies of the AMGs that were adjacently positioned, such as was seen for LB phage contig_18935 (Figure 3B). A copy of the *crtF* gene, part of the okenone synthesis pathway of pigment production was identified in a putative lytic phage. Among the phages with AMGs of interest, three LB phage-host linkages could be made with high confidence based on CRISPR-spacer homology matches, two were predicted to infect the GSB *Chlorobium chlorochromatii* CaD3 (encoding *moeB* and *iscS*), and one infecting *Pararheinheimera soli* BD- d46 (encoding *nrnA*). From the lower confidence matches (100% identity, 18-20 nucleotide coverage, and <2 mismatches), we identified nine LB phage-host pairings among the phages with AMGs of interest. This included the *crtF*-containing LB phage (contig_6928) predicted to infect the PSB *Thiocystis violascens* DSM 198, a lysogenic phage with two copies of *cysH* (contig_11073) predicted to infect the GSB *Chlorobium phaeobacteroides* DSM 266, and a phage with *thiF* (contig_43205) infecting the PSB *Arsukibacterium* sp. MJ3.

## Discussion

Here we identified viral genomes recovered from LB and PL metagenomes that include novel lineages infecting GSB and PSB, as evidenced by the long branch lengths during phylogenomic analysis (Supplementary Figure 4). Many of these novel phage lineages include phages with AMGs with potential to modify hosts’ metabolism and ecology. From this preliminary work on two proxy lakes in the Pacific Northwest, we propose that bacteriophages have the potential to affect the distribution and the energetic metabolism of GSB and PSB by modulating (I) light harvesting molecule production, (II) carbon fixation, and (III) sulfur metabolism (Figure 4).

**Figure 4.**
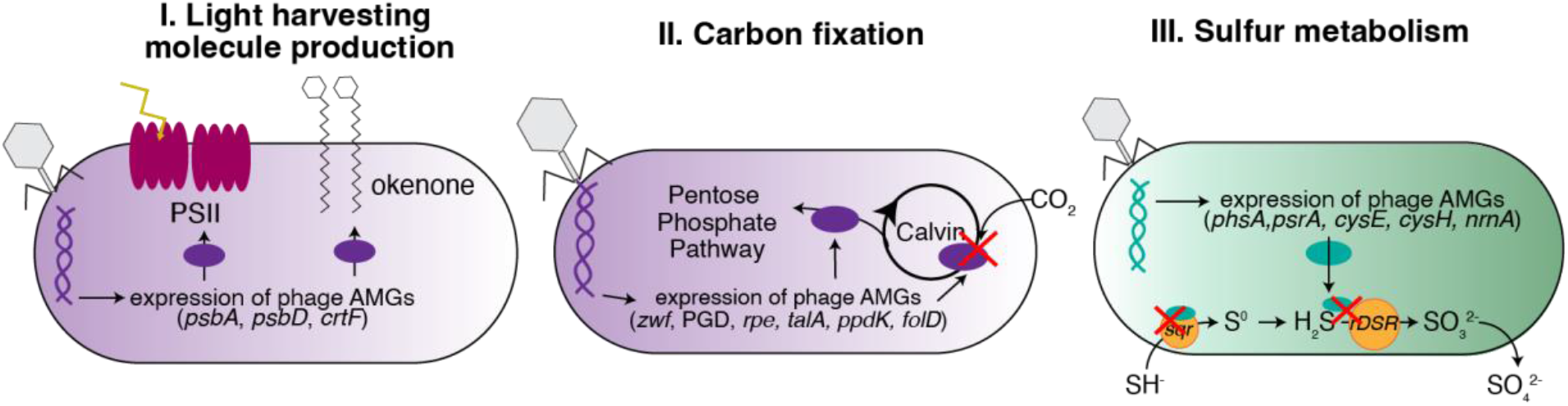
Conceptual hypotheses for viral infection influence on PSB and GSB communities. Viral predation and gene transfer affect the biosignatures of PSB and GSB by modulating their I. pigment production; III. carbon fixation; and III. sulfur metabolism.

### Viruses encode pigment and reaction center genes

Ecological distributions of GSB and PSB in euxinic water bodies and the presence or preservation of their light harvesting biomarkers (i.e., chlorobactene, isorenieratene, okenone, and their diagenetic products) in the fossil record as proxies of photic zone euxinia (Brocks *et al.*, 2005; Brocks and Schaeffer, 2008) are not fully explained by physical and chemical bottom-up controls. Our metagenomes from LB and PL presented an alternative explanation for this observation, as we identified a putative viral genome encoding a gene for the second-to-last step in okenone synthesis (*crtF*) (Vogl and Bryant, 2011) and predicted to infect the PSB *T. violascens* DSM 198 (Figure 3B). We hypothesize that viral-encoded auxiliary metabolic genes may increase the production of okenone by PSB in LB, despite the dominance of GSB in this lake. If true, this work will have important implications for interpretation of the deep time record of the diagenetic products of pigment biomarkers. This hypothesis is consistent with previous work showing that horizontal gene transfer in Lake Banyoles (Spain) results in the unexpected synthesis of BChl e and isorenieratene by *Chlorobium luteolum*, a bacterium that usually synthesizes BChl c (Llorens-Marès *et al.*, 2017). This gene transfer event offered fitness advantage to *C. luteolum* over green-coloured GSB by expansion of photo-adaptation range in a deep basin. GSB genomes have high signatures of horizontal gene transfer, reaching 24% of all genes in *Chlorobaculum tepidum* (Nakamura *et al.*, 2004). We also identified putative viral genome fragments carrying genes encoding the D1 and D2 subunits of the PSII (*psbA* and *psbD*) By modifying light reaction rates through the expression of these genes, viral infection could indirectly affect the metabolism of pigment molecules associated with reaction centers.

### Viruses encode carbon fixation genes

PSB and GSB are the main autotrophs in euxinic lakes, yet their abundances are unreliable proxies for photosynthetic activity, as demonstrated in Lake Cadagno, Switzerland (Musat *et al.*, 2008). In one growing season, the PSB *C. okenii* accounted for only 0.3% of cell abundance and 70% of the carbon uptake. In subsequent growing seasons, GSB became dominant, representing 95% of the community, but the PSB *T. syntrophicum* was responsible for 25.9% of carbon fixation (Storelli *et al.*, 2013). Given that PSB are more depleted in ^13^C than GSB using the same carbon source, their carbon isotope fractionation patterns can be used to determine the relative contributions of PSB and GSB to photosynthetic production (Posth *et al.*, 2017). During a spring bloom in Lake Cadagno three PSB strains contributed 38% to the bulk isotope signal while a single GSB contributed 62%. By fall, these PSB strains contributed 55% to the bulk isotope signal, while a single GSB contributed 45% (Posth *et al.*, 2017). Seasonal changes in the relative contribution of PSB and GSB activity to carbon isotope composition were positively correlated with cell counts (GSB were dominant in October) but had unexplained relationships with pigment concentration (Bhcl a increased and Bchl e decreased). Viral infections that increase rates of light reactions of photosynthesis while lowering carbon fixation by inhibiting the Calvin Cycle, as observed in cyanophages, could explain this pattern (Figure 4, Thompson et al., 2011).

The reductive pentose phosphate and reverse tricarboxylic acid cycle pathways utilized for carbon fixation in PSB and GSB, respectively, are shared with Cyanobacteria (Sirevåg, 1995; Tabita, 1995). Phage infections of Cyanobacterial hosts alter light reactions, the Calvin Cycle, the PPP and nucleotide biosynthesis through the expression of AMGs (e.g., *rpi, talC, tkt* and *can*; Breitbart et al., 2018). Viral infections can shut down carbon fixation while maintaining or even supplementing light reactions to support phage replication (Sullivan *et al.*, 2010; Philosof *et al.*, 2011; Thompson *et al.*, 2011; Puxty *et al.*, 2016). Therefore, viral modification of host photosynthetic machinery could be the source of unexplained patterns of carbon isotope fractionation observed in PSB and GSB. Our metagenomic analysis of LB and PL identified viral AMGs capable of interfering with the Calvin Cycle (CP12) and PPP (PGD, G6PD, *tal*; Figure 3). These observations suggest that carbon fixation rates, and therefore carbon isotope fractionation by GSB and PSB, can be modified by viral infection similar to the phage modification of Cyanobacterial carbon metabolism.

### Viruses encode sulfur cycling genes

Phototrophic sulfur bacteria oxidize inorganic sulfur compounds under anaerobic conditions. All phototrophic *Chromatiaceae* and most *Ectothiorhodospira* and GSB oxidize sulfide and elemental sulfur to sulfate, using them as electron donors for photosynthesis (Frigaard and Dahl, 2009). In our metagenomic survey of LB and PL, we found phage genomes encoding at least nine genes involved in sulfur metabolism and relay system, including genes involved in sulfur assimilation as cysteine (*cysH*, *mec*) and genes involved in the synthesis of molybdopterin, a cofactor in sulfite reduction.

We hypothesize that these viral genes deviate sulfur from the bacterial energetic metabolism towards amino acid synthesis for viral particle production. If true, viruses have the potential to modify environmental sulfur isotopic fractionation that is used in the interpretation of sulfur cycling in the geologic record. This is because the combined effects of sulfide oxidation, sulfate reduction and disproportionation influence the apparent fractionation between sulfate and sulfide isotopes (Brabec et al., 2012; Zerkle et al., 2012; Pellerin et al., 2015; Findlay et al., 2019). In euxinic lakes, isotopes of elemental sulfur are expected to correlate with photosynthetic activity (sulfide consumption) and sulfate reduction (sulfide production) (Hamilton *et al.*, 2014). Viruses encoding genes that deviate sulfur from energetic metabolism towards viral particle production may significantly modify the apparent sulfur fractionation. The presence of the gene *cysE* (serine biosynthesis) in putative phage genomes predicted to infect PSB in LB supports this hypothesis (Figure 3).

Additionally, several putative phage genomes encoding *cysH* in LB and PL may affect assimilatory sulfate reduction. *CysH* encodes a reductase that catalyzes the conversion of phosphoadenosine phosphosulfate to sulfite and is repressed under photoautotrophic growth using hydrogen sulfide as electron donor and derepressed under conditions of sulfate deficiency in PSB (Haverkamp and Schwenn, 1999). Phage alteration of this fine enzymatic regulation is a potential source of deviations in sulfur isotope fractionation.

## Conclusion

Here we describe hundreds of novel putative viral genomes from modern euxinic lakes that are analogs of early Earth oceans. We identified widespread PSB and GSB phage infections with the potential to regulate pigment production, photosynthesis, carbon fixation and sulfur metabolism, suggesting that these viruses can affect host physiology and ecology. Our preliminary observations impact the interpretation of paleoecology, paleochemistry and paleosedimentation based on biological signatures of PSB and GSB in the geologic record.

## Availability of data and materials

The Nanopore metagenomic sequencing data generated for this study for Lime Blue sediment (SRS13178833) and Poison Lake water (SRS13178834) is available in the Sequence Reads Archives (SRA) repository, under the BioProject PRJNA842402.

## Acknowledgements

We thank Molly O’Beirne for comments and discussions about the decoupling between PSB and GSB activity, abundance and biomarkers.

## Authors contributions

A. Bosco-Santos and C.B. Silveira designed the study. A. Bosco-Santos, W. Gilhooly and J. Werne collected samples and extracted DNA. S. Garcia and C. B. Silveira sequenced samples. P.J. Hesketh-Best performed bioinformatics analyses and data visualisation. First draft was written by A. Bosco-Santos, P.J. Hesketh-Best, and C.B. Silveira. All authors contributed to editing the manuscript.

## Supplementary data

**Supplementary Figure 1.**
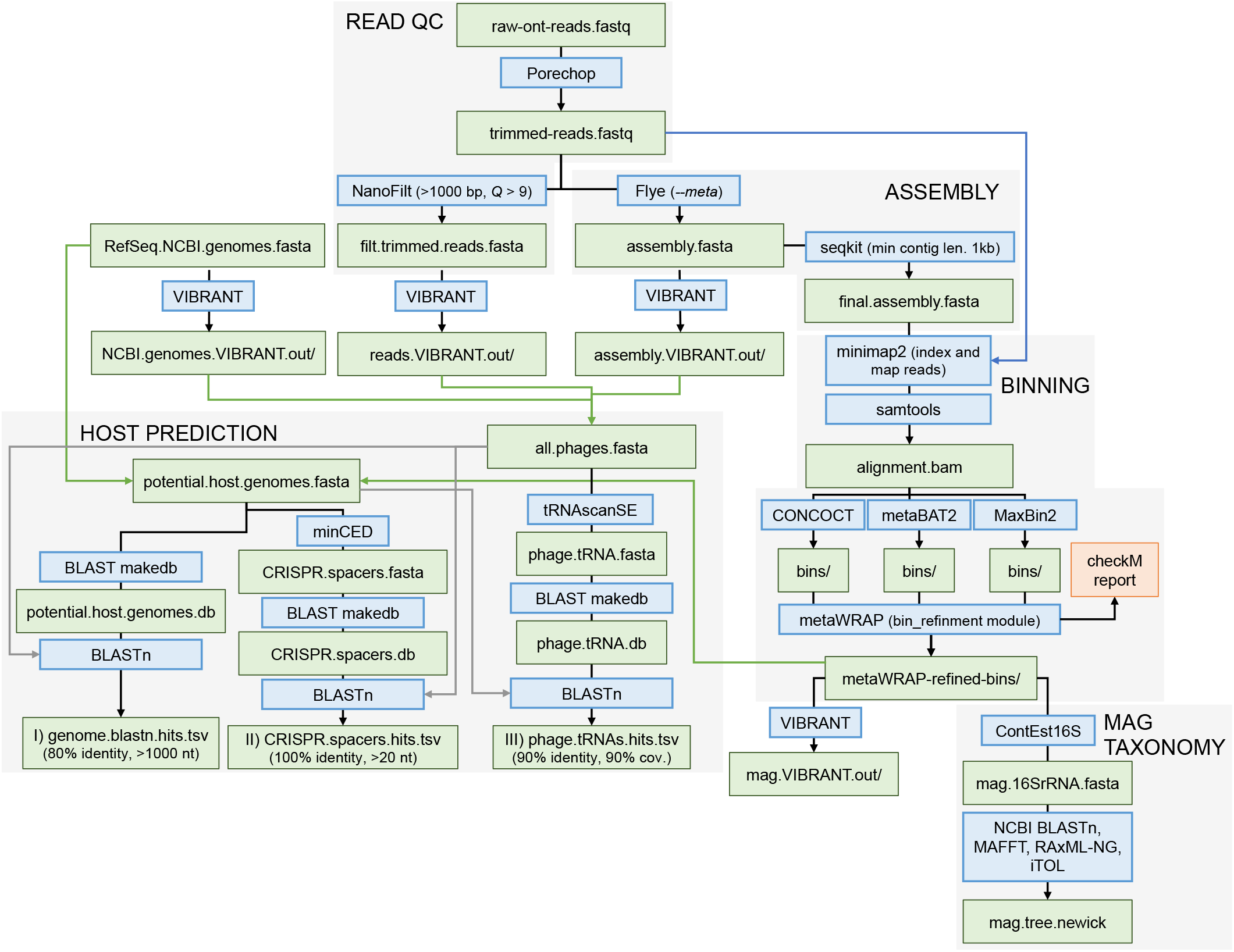
Workflow outline of metagenomic analysis conducted as part of this study.

**Supplementary Figure 2.**
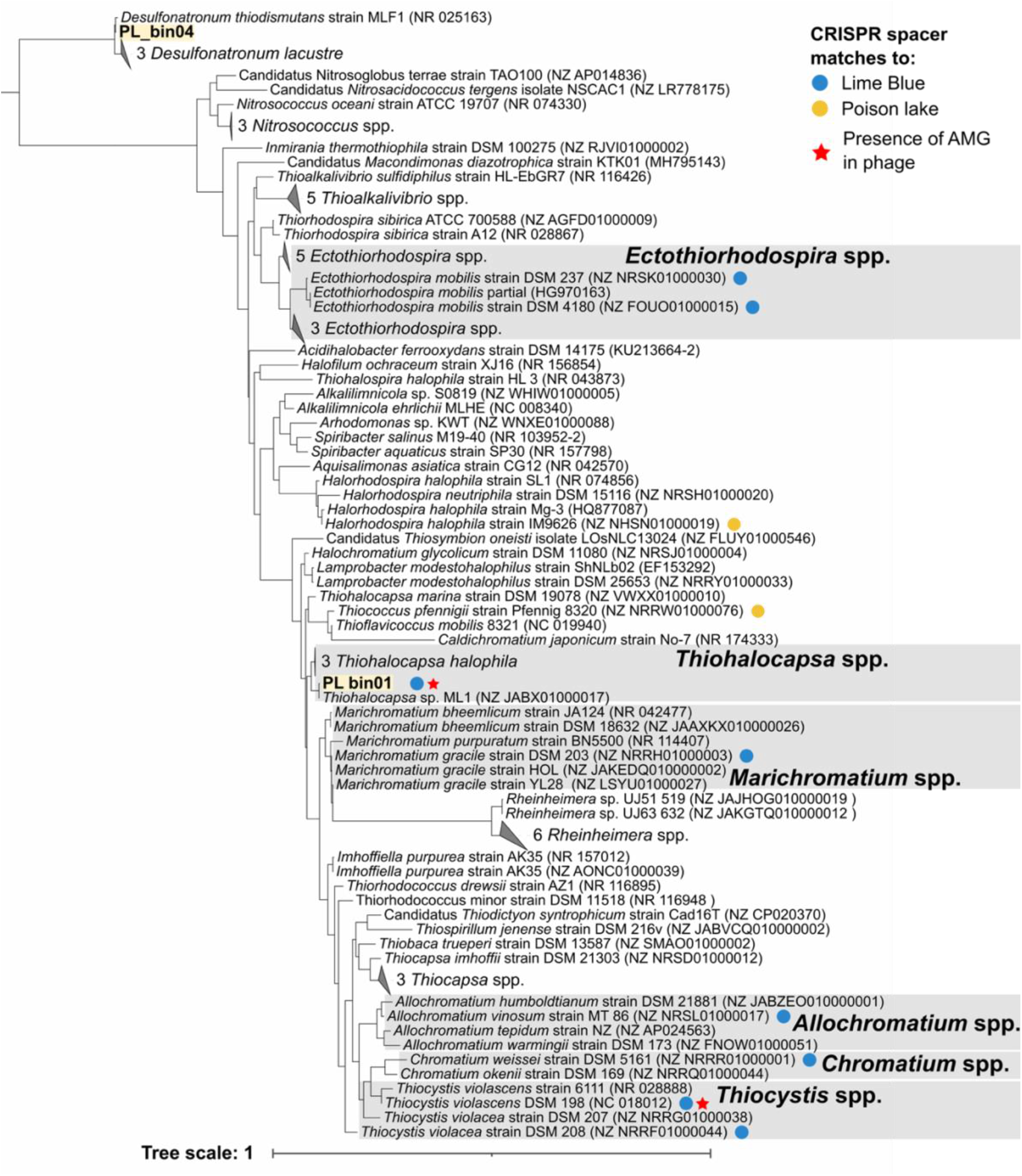
Phylogenetic tree of phages, and MAG phylogeny displaying phage hosts from the PSB as predicted by CRISPR spacer matches (minimum length 20 nt; 100% identity, maximum of 2 mismatches/gaps).

**Supplementary Figure 3.**
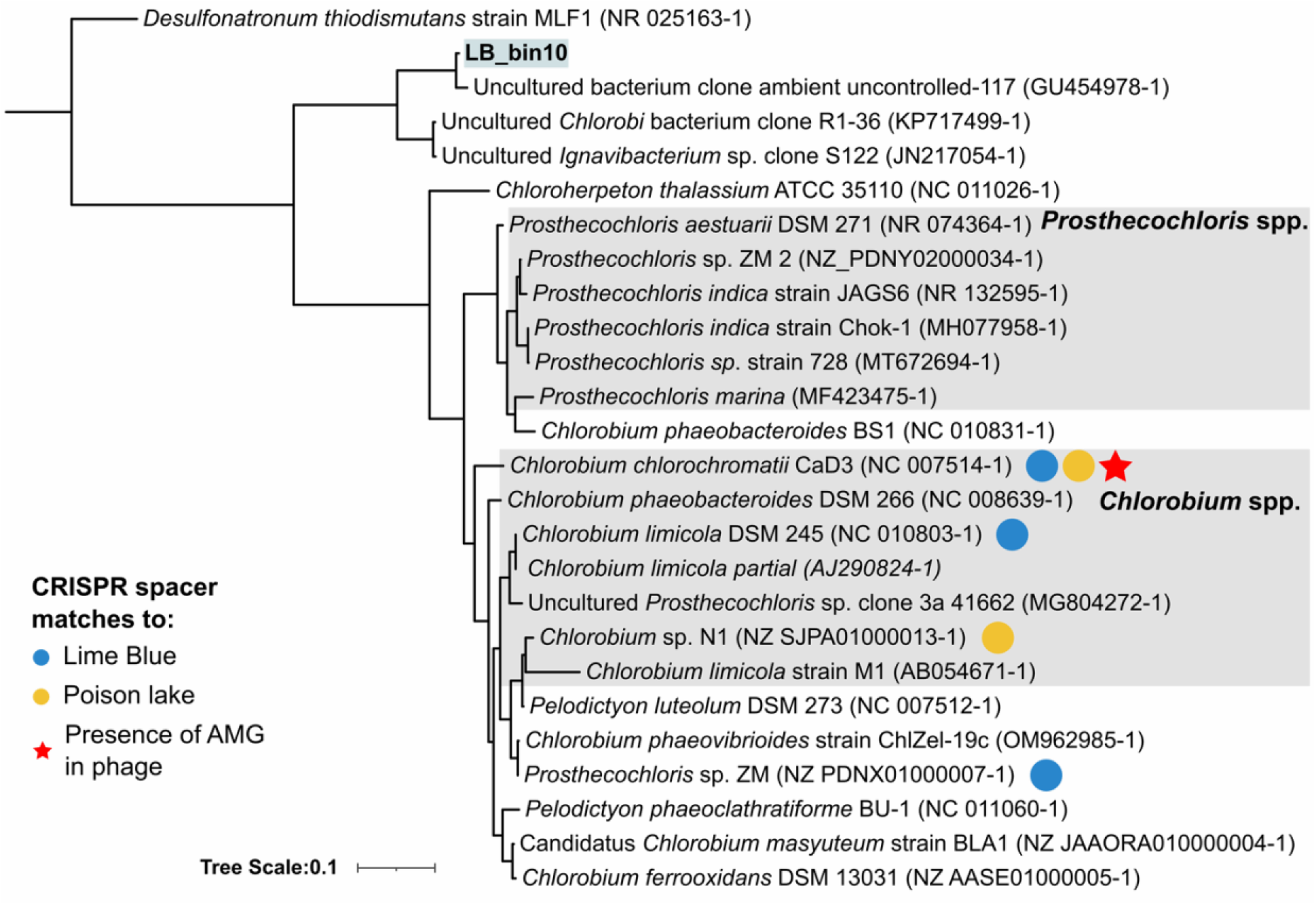
Phylogenetic tree of phages, and MAG phylogeny displaying phage hosts from the GSB as predicted by CRISPR spacer matches (minimum length 20 nt; 100% identity, maximum of 2 mismatches/gaps).

**Supplementary Figure 4.**
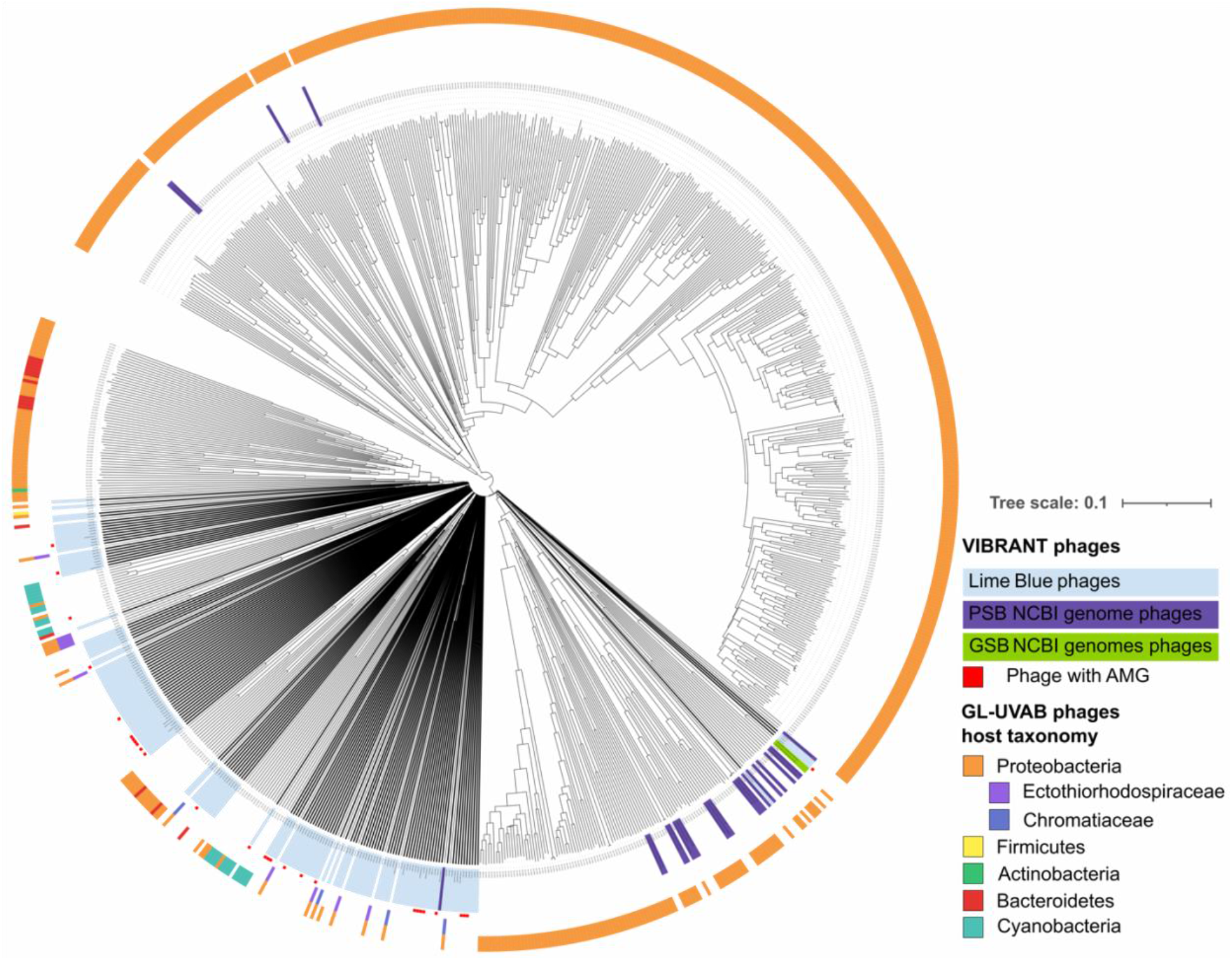
Clustering of VIBRANT identified phages from LB metagenome and PSB genomes GSB genomes and the reference phage genomes based on their Dice distance with supporting branch lengths.

**Supplementary Table 1.** Summary of purple and green sulfur bacteria genomes retrieved from the National Centre for Biotechnology Informations (NCBI).

**Supplementary Table 2.** Complete results of the predicted phage hosts pairings by sequence homology marches of complete genomes, CRISPR-spacers and tRNA sequences.

